# Affinity-mass spectrometric technologies for quantitative proteomics in biological fluids

**DOI:** 10.1101/114751

**Authors:** Huiyan Li, Robert Popp, Christoph H. Borchers

## Abstract

Proteins are the functional molecules in organisms and are therefore excellent biomarker candidates for a diversity of diseases. Immunoassays and mass spectrometry (MS) are two major technologies being used in proteomics; however, they either lack specificity or sensitivity. An emerging trend is to combine immunoassays with MS (which we call “affinity-MS”). This is an important milestone in quantitative proteomics, making it possible to measure low-abundance proteins with high specificity. The targeted enrichment and the assignment of mass-to-charge ratios to different molecules provide two selection criteria, making affinity-MS highly specific. Picogram-per-milliliter limits of detection have been obtained for many proteins. Furthermore, multiplexing capacity of >150 proteins has been achieved. This article reviews different formats of affinity-enrichment methods, and demonstrates how they are interfaced with both electrospray ionization (ESI) and matrix-assisted laser desorption/ionization (MALDI) MS. The pros and cons of these techniques are compared, and future prospectives are discussed.

## 1. Introduction

Proteins are the nanomachines in organisms and can provide direct implications on biological activities at functional levels; therefore they are excellent putative biomarkers. Protein quantification has become an important tool in disease diagnosis, prognosis, and treatment. While many clinically relevant proteins exist in the μg/mL range in human plasma, most disease-related proteins and protein biomarkers fall in the low ng/mL or pg/mL range [1]. Two major analytical technologies involved in quantitative proteomics today are immunoassays and mass spectrometry (MS), but both have their drawbacks. Combining the advantages of affinity-capture with MS detection, affinity-mass spectrometric (affinity-MS) technology is an important tool for protein quantification, holding the promise to fulfill the research and clinical needs in proteomics.

Immunoassays, such as enzyme-linked immunosorbent assay (ELISA), are currently the gold standard for protein quantification from biological samples. Affinity binders, usually polyclonal or monoclonal antibodies are used to capture the target proteins or peptides. The assay specificity highly depends on the quality of the antibody, because colorimetric or fluorescence detection methods cannot unambiguously distinguish between the target antigen and non-specifically bound compounds. Although sandwich formats using two antibodies to detect each target have been widely applied, the resulting assays suffer from cross-reactivity between reagents, and are especially problematic for low abundance proteins [2]. In addition, because two antibodies are needed for each target, this results in longer assay development period for antibody selection and also increases the cost of the assay. Major obstacles in immunoassays and potential solutions have been previously discussed by Hoofnagle et al [3].

MS directly detects the mass of the target proteins or peptides by mass-to-charge (m/z) ratio and tandem MS (MS/MS) allows sequencing of the peptides, thus offering excellent assay specificity. Peptides are commonly used as surrogates of the target native proteins for quantification due to their improved solubility and better MS sensitivity and mass resolution, with the underlying assumption that the peptide concentrations can represent -- or at least be correlated with -- the amount of the parent protein in the samples. Stable isotope standard (SIS) peptides, with the same amino acid sequences as the target peptide, but with predictable and detectable mass shifts in the mass spectra, are spiked into the sample as internal standards for absolute peptide quantification. However, with MS, a large amount of disease-related proteins at low ng/mL levels or lower in plasma cannot be detected without pre-enrichment, partly due to ion suppression from high-abundance proteins in the samples [4].

To combine the advantages of immunoassays and MS, affinity-MS technologies were developed and pioneered by Hutchens and Yip in 1993 [5]. The general procedure of a bottom-up affinity-MS assay includes selecting a predicted proteotypic peptide with suitable mass (> 800 Da) for MS analysis, producing an antibody against the peptide and immobilizing the antibody on a surface, incubating the digested sample and SIS peptides with the antibody for immuno-enrichment, and measuring the endogenous (END) and SIS peptides with a mass spectrometer. The immuno-enrichment step concentrates and purifies the target molecules from samples, therefore improving assay sensitivity compared to direct MS analysis without enrichment. Only one antibody is required in affinity-MS to enrich target proteins or peptides, instead of two antibodies as in a sandwich ELISA or western blot, significantly reducing the time and cost for assay development. The second round of selection is performed by the mass spectrometer with ultra-high specificity, eliminating or at least reducing the non-specificity issues of immunoassays and saving time and the cost of screening antibody pairs. In top-down affinity-MS, intact proteins are enriched and detected, making this technique capable of distinguishing protein isoforms at low concentrations which cannot be discriminated with immunoassays, as long as they have different molecular ions [6].

In this article, we will review recent advances in the development of affinity-MS assays and evaluate their applications for quantitative proteomics. Different immuno-enrichment strategies and their interfaces with either electrospray ionization (ESI) or matrix-assisted laser desorption/ionization (MALDI)-MS are compared in terms of assay performance. Current limitations and future perspectives will be discussed.

## 2. Affinity-mass spectrometric technologies

### 2.1 Affinity-enrichment on beads

Microbeads have become the most commonly used tools for immunoprecipitation (IP). In IP, agarose or superparamagnetic beads are functionalized with antibodies to capture proteins or peptides. Different methods for binding the antibody to the beads have been discussed previously [7]. Compared to affinity column-based methods, microbead handling is much easier and potentially more cost effective. Moreover, microscale beads have a high surface-to-volume ratio, therefore increasing antibody binding capacity and thus improving IP efficiency over column-based affinity enrichment techniques. Small sample volumes (down to 2 μL) can be enough for IP with the beads [8]. In affinity-MS approaches, bead-based enrichment of target proteins and peptides has been interfaced with either MALDI- or ESI-based instruments, as described below.

#### 2.1.1 Immuno-MALDI (iMALDI)

iMALDI combines enrichment of target peptides using antibodies bound to affinity-beads. Immediately after the immuno-enrichment and subsequent washes, the beads are directly spotted on a MALDI plate. The antibody-peptide complex is disrupted on-target by the application of acidic MALDI matrix.

iMALDI technology was first reported a decade ago by Borchers *et al.* [9]. Protein digests, together with the SIS peptides, were incubated with antibody functionalized agarose beads (40-165 μm in diameter) for immuno-enrichment. Notably, a single bead, with the capability of binding ~ 12 fmol protein or peptide per antibody-coated bead, was placed on the target plate after immuno-enrichment. 0.3–0.5 μL of matrix solution was then added and dried, resulting in a ~ 1-2 mm spot. A LOD of ~ 12 fmol was achieved with the single bead assay. In the future, the assay LOD can be further improved by delivering a much smaller volume of matrix solution and concentrating the target peptides in microscale spots for MALDI analysis.

To date, iMALDI technology has been applied to biomarker detection for a variety of diseases. Jiang et al. developed a tandem iMALDI diagnostic assay to quantify the epidermal growth factor receptor (EGFR) protein in breast tumor cell lysate and biopsy samples [10]. Anti-peptide antibody was bound covalently on CNBr-activated sepharose beads, and incubated with the protein digest and SIS EGFR peptide for 2-4 h. One attomole of the target peptide could be detected in buffer, and the protein was detectable in cell lysate from only a few cancer cells and 1/25 of the biopsy samples. For the diagnosis of *F. tularensis,* the IglC protein in *F. tularensis* bacteria was targeted. Nasal swab samples from mice inoculated with *F. tularensis* were collected, digested, and spiked into buffer or human plasma [11]. The iMALDI method achieved LODs of 14 amol of synthetic peptide enriched from buffer, 8 CFU bacteria from a bacterial digest in buffer, 69 amol synthetic peptide spiked into plasma, and 800 CFU of bacteria spiked into human plasma, which is comparable to most ELISA methods, but with higher specificity due to MS/MS capability.

Another successful application of the iMALDI technology has been the quantification of Angiotensin I (Ang I), an octapeptide that is quantified to determine a patient’s plasma renin activity (PRA) for the diagnosis of primary aldosteronism [12, 13]. Radioimmunoassay and ELISA assays have been widely used in the clinical measurement of PRA, but suffer from cross-reactivity, leading to false positive signals. In the iMALDI PRA assay, a commercially available anti-Ang I antibody was bound to magnetic Protein G Dynabeads, and incubated with the human plasma samples, together with the spiked Ang I SIS peptide (see Fig. 1a). The LOD and linearity obtained by iMALDI fell within the physiological ranges, and the inter- and intra-day coefficient of variation (CV) was ~ 17% when the assays were conducted manually. Later, a duplexed assay for the detection of both Ang I and Ang II was developed and an LOD of ~ 10 pg/mL was achieved for both peptides [14]. Recently, the assay has been automated using a liquid handling system which achieved CVs below 10% [15]. The method correlated well with a clinically used LC-MS/MS assay when measuring clinical patient samples, and had a 7.5-fold faster analysis time than LC-MS/MS, allowing for the analysis of up to 744 samples per day. More recently, a convenient-to-use method has been developed in our lab that simplifies the iMALDI sample preparation and avoids the use of the complex liquid handling system, making iMALDI accessible to most research and clinical laboratories [Li, *et al.* Submitted].

**Fig. 1.**
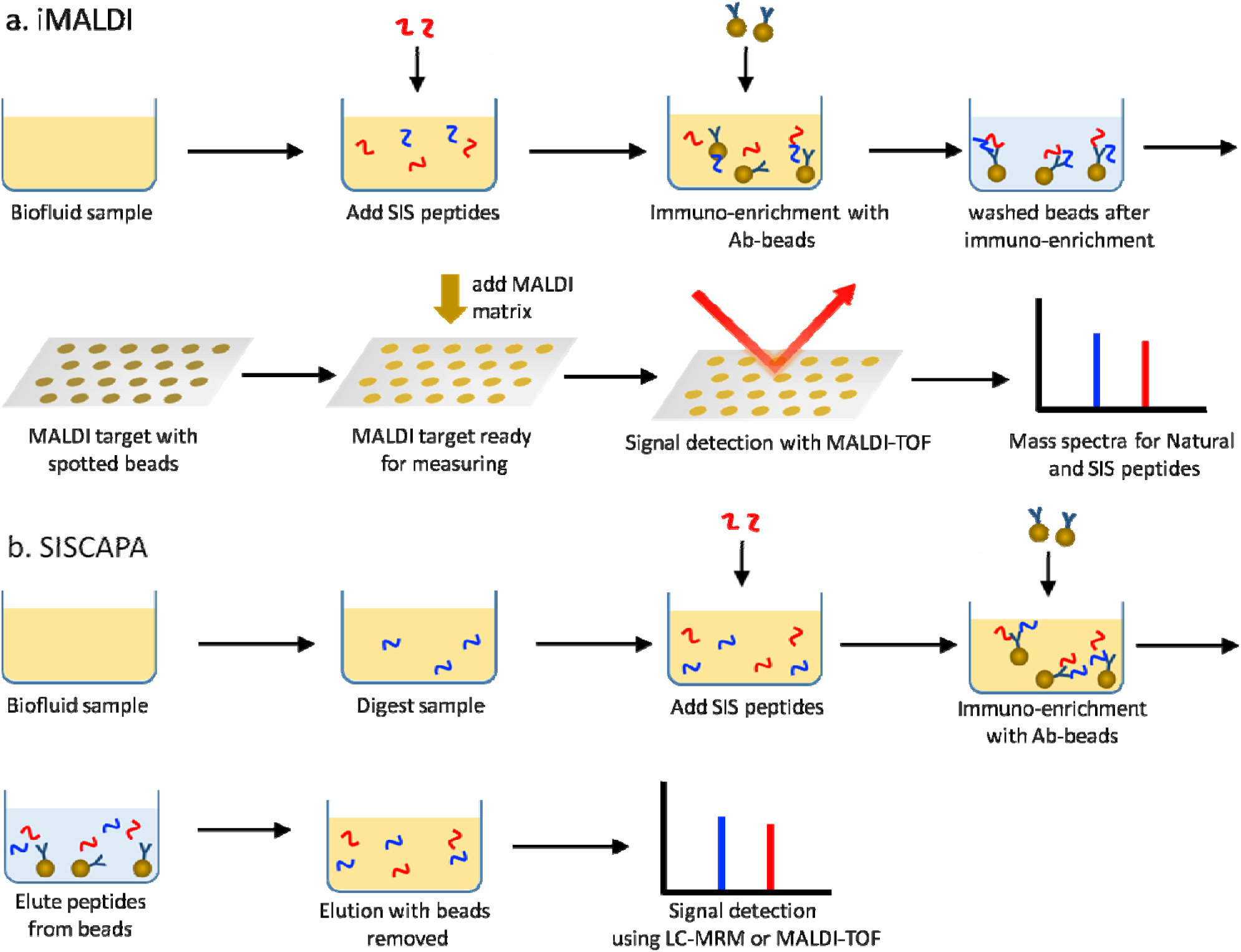
Process flow for (a) iMALDI and (b) SISCAPA. Both methods involve co-capture of analyte and SIS peptides with antibody-coated beads. In iMALDI, beads are directly transferred onto a MALDI target plate after immuno-enrichment. In SISCAPA, the captured peptides are first eluted from the beads, followed by measurement using LC-MRM or MALDI-TOF.

The advantages of iMALDI are the elution of target molecules on the MALDI plate, thereby reducing sample losses, e.g. due to plastic adsorption. Another advantage is the high-throughput of MALDI-TOF, which allows analysis times of typically <10 seconds per sample. Furthermore, even though MALDI-TOF is known for highly variable signal intensities, e.g. due to variable ionization efficiency from spot to spot, the use of SIS peptides compensates for these intensity fluctuations and allows highly precise quantitation results. Another advantage is that the detection limits appear to be lower than assays where the analytes are eluted from the beads, and then spotted on the target as a separate step, such as the techniques described below. This probably because spotting prior to elution avoids the loss of analyte to the walls of the tube that contains the beads during the elution step.

#### 2.1.2 immunoMRM (iMRM)

iMRM also called SISCAPA was proposed by Anderson et al [16]. In the initial work, anti-peptide antibodies were covalently immobilized on columns, and the target peptides together with SIS standards were enriched. After elution from the column, the peptides were quantified by multiple reaction monitoring (MRM). Later, a magnetic bead-based platform was developed, and the immuno-enrichment helped improve the ion signals by three orders of magnitude, capable of quantifying protein biomarkers in the ng/mL range [4]. The assay workflow is illustrated in Fig. 1b. Notably, it was found that adding a trypsin inhibitor to the samples after protein digestion could prevent the antibody from being digested by the remaining trypsin in the samples, thereby improving peptide recovery. The bead-based SISCAPA was automated with a rotary bead trap device for simplified sample preparation [17]. A KingFisher liquid handling system was then used to automate a 9-plex SISCAPA assay. The assay achieved LODs in the ng/mL range from 10 μL of plasma, and a median CV of 12.6%. The LOD was extended to low pg/mL levels when 1 mL of plasma was used [18].

To demonstrate the transferability of the SISCAPA-MRM approach between labs, and to evaluate the sources and degree of lab-to-lab variation, a four-phase inter-laboratory study was conducted by performing an automated 8-plex SISCAPA assay in three independent labs [19]. Samples were prepared at one site to avoid the variations from initial sample preparation. The standard curves showed a comparable linear response between assays and laboratories in the concentration range of 0.23-167 fmol/μL for all the peptides, implying that the assay can be used for relative protein quantification between samples without running a preliminary assay for the estimation of concentration. LODs of 1 ng/mL were achieved from 30 μL plasma, and the median intra- and inter-laboratory CVs above LOQ were 11% and <14%, respectively. It was found that tryptic digestion contributed most of the assay variability and reduced the recovery of the peptides from the proteins. Interestingly, the use of heavy protein standards rather than SIS peptides on average doubled the assay accuracy, and improved the assay precision by 5% [19]. Overall it was demonstrated that immuno-MRM assays can be reproducible across laboratories.

Recently, automation of the tryptic digestion and the immuno-enrichment steps has been developed for SISCAPA, eliminating the sample cleanup after digestion and simplifying sample preparation. The assay reproducibility has been significantly improved, with an intra-assay CV of 3.4% and inter-assay CV of 4.3% [20].

Next, several technology advancements have been reported to improve the SISCAPA assay throughput and multiplexing capability. Chromatography-free SISCAPA was combined with solid-phase extraction (SPE)-MS/MS for sensitive and rapid MRM-based quantification of peptides [21]. An ultrafast online SPE platform coupled to a mass spectrometer was used, and the time for transferring the digested sample to the mass spectrometer was only ~7s, 300-fold faster than conventional LC-MRM-MS analysis. Non-specific binding of peptides to the magnetic beads was reduced by choosing optimal beads, and by adding a final wash with 3:1 acetonitrile/PBS before elution of the peptides from the beads. Using this method, a 2-plex assay quantifying peptides from protein C inhibitor and soluble transferrin receptor was conducted in 159 sera collected from 51 prostate cancer patients [22]. Recently, a 50- plex SISCAPA assay was achieved by sequentially capturing analytes in the same sample using different types of beads for each round of capture [23].

In addition, SISCAPA has been used to quantify both modified [24, 25] and non-modified proteins. For example, 6 candidate colon cancer biomarkers were measured from plasma of colon cancer patients and healthy controls [26]. The data showed a strong correlation with results from conventional ELISA. In another study, a 69-plex assay measuring proteins in the phospho-signaling networks was developed [25]. Both phosphorylated and non-phosphorylated peptides were detected in parallel in human cells and human cancer tissues. Recently, a temporal study of proteins in dried blood spot (DBS) was performed by automatically measuring multiple proteins in 14 individuals over 6.5 years [27]. The extracted proteins from DBS were firstly digested to peptides for analysis, thus circumventing the loss of activity issue at the protein level upon drying on the collection cards. To calibrate spot-to-spot hematocrit and blood-volume variations, a panel of ‘normalization proteins’ were used. With automation, CVs of less than 5% were obtained for all the proteins, allowing the comparison of each individual's own protein levels over time. The results showed that while the variations of protein levels in healthy individuals were relatively small, a large departure from baseline values were observed due to infections, normal pregnancy, and season of the year.

Recently, SISCAPA has been interfaced with MALDI-TOF MS detection, enabling liquid-chromatography-free protein quantitation [28, 29]. SIS peptides were added into the digested samples, and captured by antibody-beads together with the endogenous target peptides. Next, the beads were washed, and captured peptides were eluted from the beads and measured with a MALDI-TOF instrumentation. With the help of SIS peptides, CVs of 1.5-3.7% were achieved on measuring different peptides, confirming the robustness of this MALDI technology[29].

### 2.2 Affinity-enrichment on a chip

In this section, recent literature describing affinity-enrichment performed on a chip will be reviewed, including surface enhanced laser desorption/ionization (SELDI) technology, surface plasmon resonance imaging (SPRI) coupled with immuno-MALDI-TOF, and mass-tag enhanced MALDI-MS.

#### 2.2.1 SELDI-TOF-MS

The first affinity-MS work based on SELDI technology was reported back in 1993 [5]. In a typical SELDI-MS assay, the affinity-enrichment of proteins on a solid surface is achieved by either chemically functionalizing the surface in an array format according to general or specific properties such as hydrophobicity or charge, or coating the surface with affinity binders such as antibodies or DNAs [30]. For example, on an ionic exchange array, charged molecules are reversibly bound to the chip [31]. Samples need to be diluted in a binding buffer specifically designed for different chips, or mixed with denaturing buffer, but usually no digestion step is involved and intact proteins are measured. Similar to MALDI, an acidic matrix solution, such as sinapinic acid in acetonitrile is applied to the sample spots. A chip reader consisting of a Laser Desorption/Ionization (LDI)-TOF-MS instrument is then used to measure the m/z value of the proteins, see Fig. 2a.

**Fig. 2.**
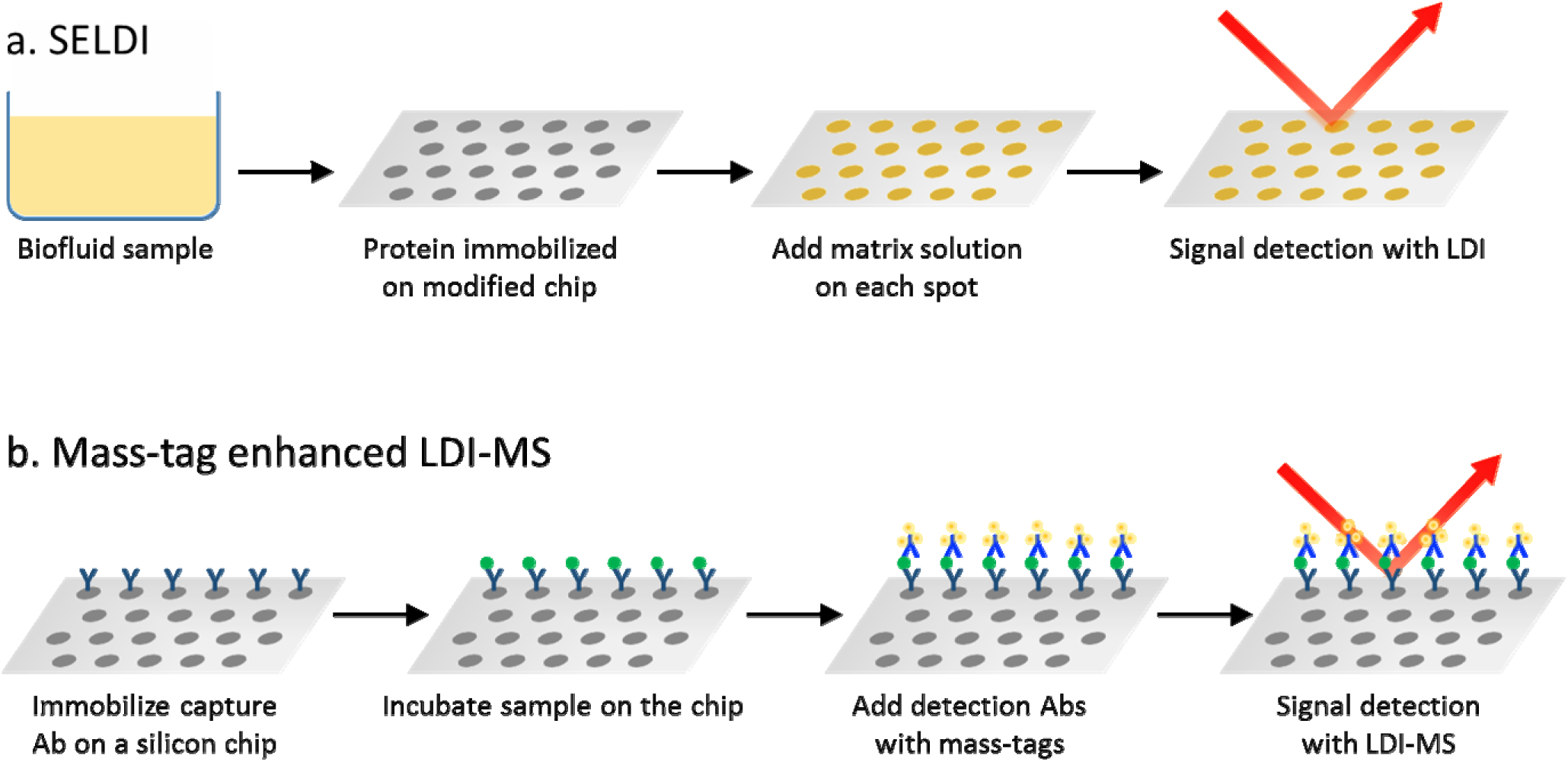
Schematic for (a) SELDI and (b) mass-tag enhanced LDI-MS. For both methods, immuno-enrichment is performed on a chip, and signals are detected by LDI-MS.

In 2002, SELDI was used to identify proteomic patterns for the diagnosis of ovarian and prostate cancer [32, 33]. However, in validation studies, the patterns found by SELDI failed to separate patients from normal controls [34, 35]. The “SELDI fiasco” was attributed to SELDI study sample bias and the differences in study design [36]. Later, Bio-Rad introduced ProteinChip^®^ SELDI System, which used a better TOF-MS instrumentation, defined more appropriate sample preparation protocols including the fractionation of complex biological samples before SELDI analysis, and optimized the sample dilution, buffer composition, and the methods for the application of matrix to the sample arrays. In addition, rigorous data collection and analysis workflow was developed for biomarker studies using the SELDI system [37].

Recently, SELDI has been used in protein quantification in a variety of diseases. Serum samples from gastric cancer patients and normal controls were measured, and five proteins were identified as the best candidate biomarkers for diagnosis [38]. One protein was recognized for distinguishing different stages of gastric cancer with high specificity and sensitivity. SELDI-TOF-MS was also combined with magnetic beads for the quantification of potential serum protein biomarkers for the diagnostics of active tuberculosis, with 35 m/z peaks detected relating to tuberculosis [39]. In addition, the use of SELDI for the profiling of saliva proteome has been reviewed by Ardito et al [31]. In most cases there are no digestion steps involved in SELDI, so most applications have been limited to relatively small proteins, and further sample processing using gels and tryptic digestion for liquid chromatography analysis was conducted [40].

A variation of SELDI-TOF is the application of antibodies to flat surfaces to enrich target molecules directly on-plate. Recent developments in ambient ion soft landing [41] have made it feasible to produce MALDI-TOF compatible surfaces with permanently deposited antibodies. Pompach *et al.* used phenotyping which has a potential clinical application in predicting complications in patients with diabetes mellitus [42]. LOD of less than 10 fmol has been achieved with 150 kD protein spots, with a CV of <11% [43].

#### 2.2.2 Surface Plasmon Resonance (SPR) coupled immuno-MALDI-TOF

To perform immuno-MALDI-TOF assays, antibodies can be immobilized on the MALDI target plate, and MALDI analysis can be conducted directly on the same chip after immuno-enrichment. This on-chip immuno-enrichment also makes it feasible to couple MALDI measurement to SPR, allowing the realtime analysis of the kinetics and thermodynamics of antibody-antigen interactions. Bellon *et al*. developed an on-chip MALDI detection assay based on a microarray SPRI biochip with a gold-covered surface [44], in which antibodies were deposited in an arrayed, and incubated with sample solutions. A key challenge of combining SPR with MS is that the sensitivity of SPR has a bias towards high molecular weight molecules and it is difficult to detect small molecules with low-abundance [44], while MS analysis usually requires digestion of proteins into peptides. To solve this problem, after SPR measurement, an on-chip digestion was performed, by dropping a solution containing trypsin onto the protein spot, followed by adding a solution of matrix for MALDI analysis. The mass spectra showed several peptides that were specific for the target proteins, as well as some fragments from trypsin autolysis and from the antibodies. A LOD of low femtomole per square millimeter was achieved [44]. Later, the process for fabricating an antibody array biochips were simplified [45], and an on-chip reduction step was included before tryptic digestion to achieve femtomole-level LODs in human plasma [46].

#### 2.2.3 Mass-tag enhanced laser/desorption ionization (LDI)-MS

Mass-tags, such as peptides or boronolectin, have been conjugated to antibodies and were firstly used in laser-based mass spectrometry for tissue imaging [47-49]. These mass tags are photo-cleavable and can be released and desorbed by the laser; each tag is specific to a different target protein and has a different mass which can then be detected. This helps avoid the spectral overlap issues in fluorescence-based imaging.

Recently, a mass-tag enhanced LDI-MS assay for the detection of intact proteins in plasma has been developed [50]. In a sandwich immunoassay targeting prostate specific antigen (PSA), the capture antibody was immobilized on a porous silicon chip, the surface was then blocked with milk and then washed. Next, samples were incubated for 30 min, and following washing, mass-tagged detection antibodies were added. To fabricate the tagged detection antibody, triphenylmethyl-(trityl) based photo-cleavable tags were coupled to avidin, and bound onto the biotin labeled detection antibodies. Signal amplification was achieved by attaching multiple tag-coupled avidin molecules onto each detection antibody, see Fig. 2b. A LOD of 200 pg/mL (60 amol per spot) PSA was obtained from 10 μL serum, comparable with conventional ELISA. CV was relatively large, depending on the efficiency of the tagging reaction. In this work, the mass-tag functioned as a secondary detection in a sandwich immunoassay. No protein digestion procedure was required and the intact protein was detected. In addition, unlike MALDI, no matrix solution was used for ionization, avoiding background signals from the matrix compound. Further, a panel of mass tags with different masses could be produced in the future, allowing for multiplexed assays. However, mass-tag based detection cannot directly provide the m/z of the analyte molecules, and the assay's specificity depends highly on the antibodies. Thus, extensive selection and validation of antibodies is needed beforehand.

### 2.3 Affinity-enrichment in 96-well plates

Affinity-enrichment has also been performed in 96-well plates. Enzyme linked immuno mass spectrometric assay (ELIMSA) combines conventional sandwich ELISA with LC, ESI, and tandem mass spectrometry for highly sensitive detection of biomolecules [51]. The small molecule products of the reporter enzymes, horseradish peroxidase (HRP) and alkaline phosphatase (AP) can be ionized efficiently and detected with a mass spectrometer, at femto to picomole amounts. ELIMSA utilizes the sandwich format of a conventional sandwich ELISA by using two antibodies to target each protein, and detects the ionized products of the reporter enzymes (HRP and AP) by mass spectrometry. Whole proteins can be detected without digestion, and the two antibodies provide a double-selection process for each protein, improving assay specificity.

Combined with the enzyme amplification from ELISA, an attomole-level LOD was achieved in the detection of PSA, which is more sensitive than radioimmunoassays and electrochemical detection methods [51]. Later, the substrate pyridoxamine-5-phosphate was used in ELIMSA to yield the enzyme product pyridoxamine, which has better chromatographic characteristics than the conventionally used HRP system [52]. A LOD of 1 pg/mL was achieved by measuring diluted PSA. In a more recent study, the alkaline phosphatase-streptavidin (AP-SA) probe was quantified by ELISMA with a linear range from 0.1 to 50 pg/mL using only 0.5 μL of plasma sample, and a LOD of 50 pg/mL was determined by spiking PSA into 100 μL plasma samples, with a CV of less than 10% [53]. Although only used in a single-plex format so far, ELIMSA has the potential to be multiplexed, but -- as for the mass-tag enhanced assay -- the specificity of ELIMSA depends on the quality of the antibody pairs, therefore requires the validation of the antibodies before the assay.

### 2.4 Affinity-enrichment in columns

Another approach that combines affinity-enrichment with MS uses columns as the solid support for the affinity reagents. A representative schematic is shown in Fig. 3. Dufield *et al.* utilized this method to enrich aggrecanase fragments followed by LC-MS/MS analysis to monitor aggrecanase activity in osteoarthritis [54]. Multiple protein G columns were prepared for these multiplexed assays. Proteins in the samples were digested and then a multidimensional (immunoaffinity/reversed-phase) LC-MS/MS analysis was performed. The immunoaffinity columns were integrated in the LC-MS/MS system, the other key components of which included a loading pump, an autosampler, on-line solid phase extraction, and a triple quadrupole for MRM analysis, see Fig. 3. An LOD of 0.5 fmol aggrecan peptides were obtained from urine samples, with intra- and inter-day CVs of less than 15%.

**Fig. 3.**
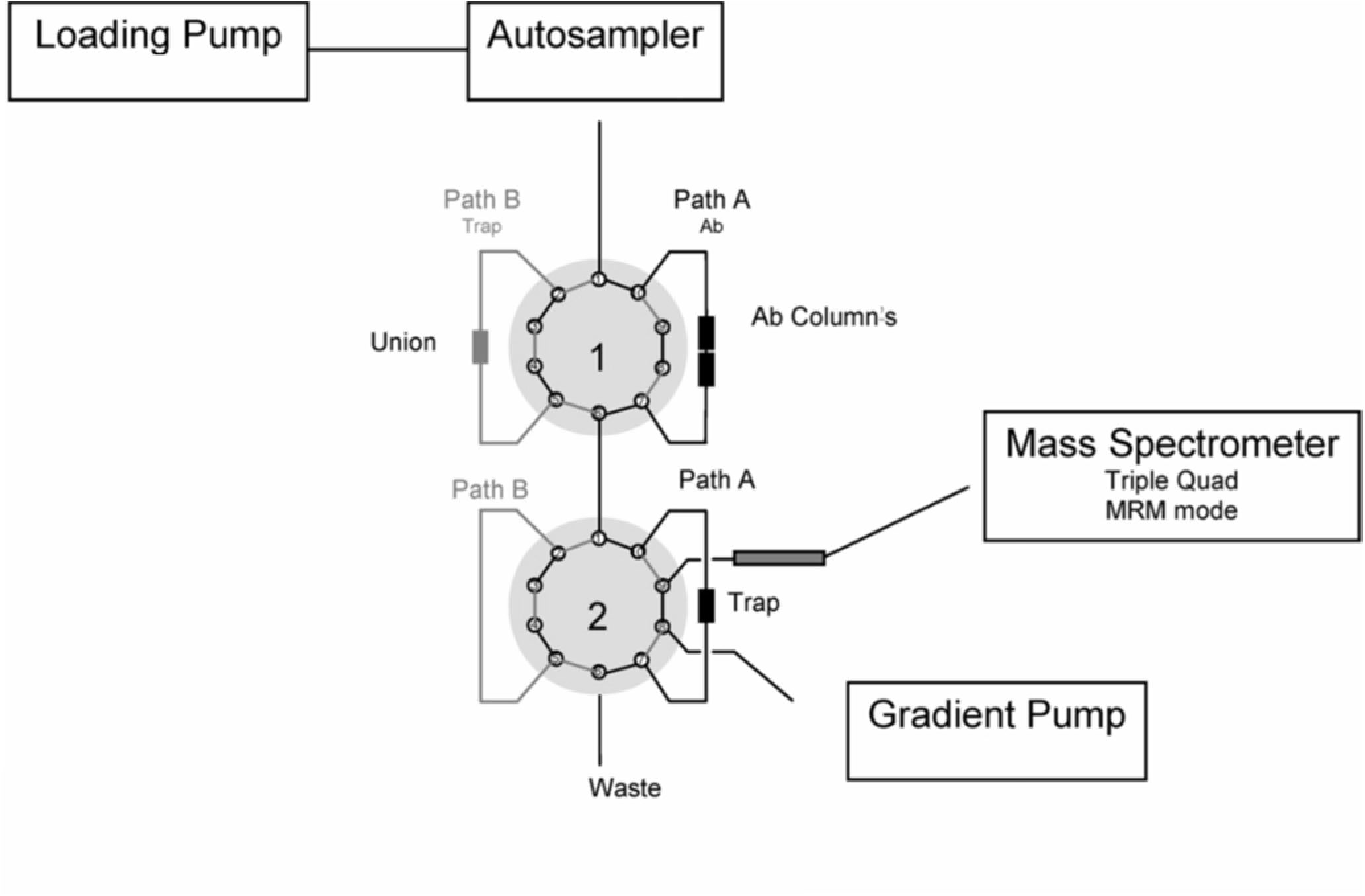
Assay procedure with affinity-enrichment steps performed in on-line columns for the detection of aggrecan fragments. Adapted from Reference [54], with permission.

In another work by Neubert et al., a high-flow on-line immuno-enrichment column was linked to nanoflow LC and selected reaction monitoring (SRM)-MS for measuring total salivary pepsin/pepsinogen [55]. The immuno-affinity column had a relative high antibody binding capacity, leading to high analyte capture efficiency. The sample injection volume ranged from low microliters to one milliliter, and the assay sensitivity was scalable with variable injection volume, meeting different requirements on the assay sensitivity without changing the configuration of the chromatography. Also, no extensive reversed-phased chromatography was required due to the simplified peptide mixtures from immuno-enrichment, reducing the chromatographic cycle time to only 15 min per sample, allowing the analysis of approximately 100 samples within 24 h. Pepsinogen as low as 4 fmol/mL was detectable in human saliva, with inter-assay CVs of 1.2-14.9% for different analyte concentrations.

These on-line immuno-affinity columns are easy to be integrated in the LC-MS system, providing an additional selection parameter for improving assay specificity and sensitivity. Another benefit of using immuno-affinity columns is that these columns might be used for hundreds to thousands of assay runs depending on the antibody, improving reproducibility and assay robustness [55]. However, as the immuno-affinity columns are integrated on-line, some samples, such as highly viscous ones, might not be well compatible with the chromatography systems.

### 2.5 Affinity-enrichment in tips

Immuno-enrichment has also been conducted in a tip-format. The pipette tips are essentially microcolumns which are packed with a material functionalized with specific ligand-binding reagents such as antibodies. The concept of the mass spectrometric immunoassay (MSIA) was firstly presented by Nelson et al in 1995 [56]. To date, MSIA has been successfully commercialized and automated for high-throughput quantification of picogram levels of analytes from complex samples. As shown in Fig. 4, in a typical MSIA assay, antibodies are immobilized in the porous pipette tips, and 100-1000 aspiration-dispense cycles allow the enrichment of target proteins from samples. Next, the tips are rinsed, and the captured proteins are eluted and measured by either MALDI or ESI-MS. Thus far, most MSIA assays have been combined with top-down MS, and they have been extensively used for quantifying different protein isoforms, which might be not distinguishable if bottom-up MS or ELISA-based assays had been used, especially when these isoforms were not known in the literature.

**Fig. 4.**
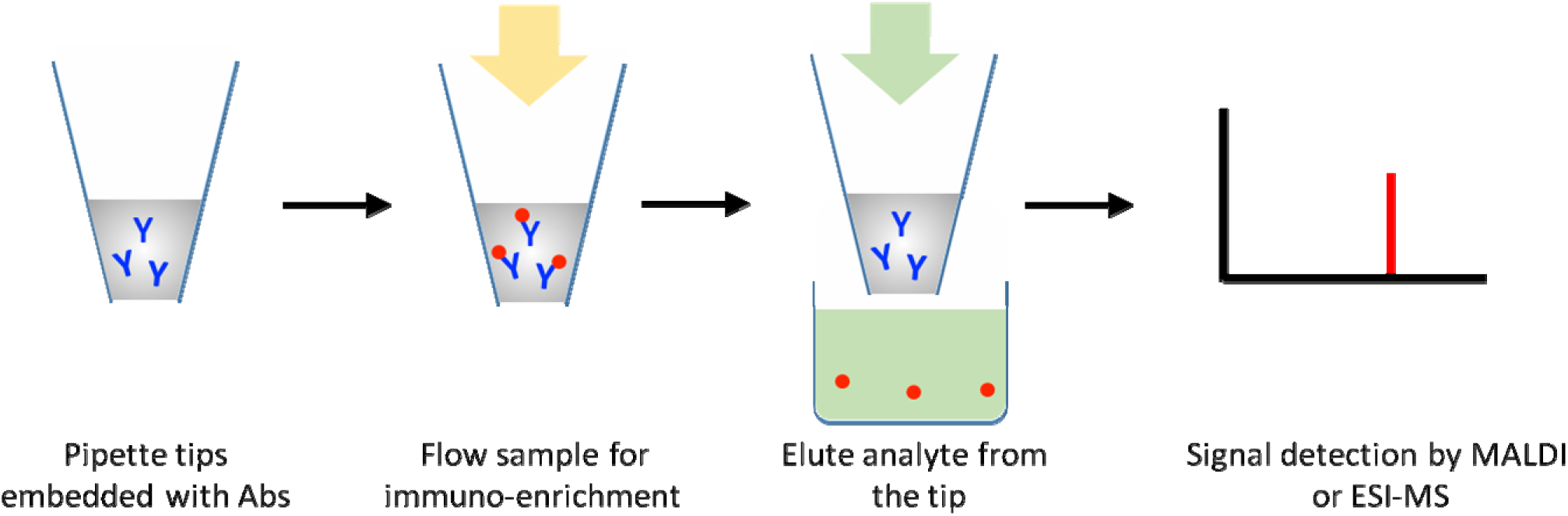
Assay schematic with affinity-enrichment in pipette tips. Following elution, either MALDI or ESI-MS is used for signal acquisition.

For detecting intact insulin and its variants, anti-human insulin antibodies were immobilized onto commercial MSIA tips [57]. Insulin metabolites and synthetic insulin analogs were measured in human plasma. A LOD of 1 pM concentration was achieved, with an inter-day CV of less than 8%. Importantly, isoforms that were not known to exist in plasma were detected in diabetics. A robotic pipette station was used to measure one 96-well plate in only 2 hours.

In another work, the longitudinal changes in 4 proteins -- including their native forms, 13 variants with post-translational modifications (PTMs) and 7 single nucleotide polymorphism-derived variants -- were quantified in 500 healthy individuals with MSIA-MALDI-TOF assays [6]. The MALDI matrix solution was used to elute the proteins from the tips, and the eluates were directly delivered onto a MALDI target. This constituted the first large-scale study of protein variants and their longitudinal changes over a six-month duration. In another example, a 3-plex MSIA assay was achieved by immobilizing a mixture of antibodies in a tip. The antibody-coated tips were found to be stable for more than 6 months at 4°C [58]. Recently, MSIA was coupled to LC-MS for measuring therapeutic antibodies. Interestingly, after immuno-enrichment, the analyte antibodies were denatured, reduced, alkylated, and digested on the tip before elution [59]. Recent development in MSIA for the characterization of proteoforms have been recently reviewed by Nedelkov et al [60].

To further improve assay throughput, a simplified sample preparation, two pipetting robots and a high-throughput MALDI-TOF was used to measure insulin-like growth factor 1 (IGF1) [61]. 1054 samples were measured within ~9 h, with intra- and inter-day CV of less than 10%. This is probably the highest assay throughput reported to date. However, the assay may take longer if protein digestion steps are required.

Although mostly used as a top-down approach, MSIA has the potential to be used for quantifying digested proteins, if anti-peptide antibodies are immobilized in the tips. In a recent work, immuno-enrichment has been performed in tips using an automated system to achieve multiplexing levels of up to 172, with a median LOD for all peptides of 71.5 amol/μL and CV of less than 15% at the LOQ [62]. 96 syringes, each equipped with a Protein G cartridge for the enrichment of target peptides, were fitted onto the Agilent AssayMAP Bravo microchromatography platform. After the screening of 172 antibodies cross-linked in cartridges, a 110-plex assay was performed in plasma samples from patients with cardiovascular diseases. This constitutes the most highly multiplexed affinity-MS assay to date. Notably, the antibody binding capacity of the cartridge was 275 μg IgG per cartridge. Considering that only 1-2 μg of antibodies are needed per target, a multiplexing level of 140-300 could theoretically be achieved. However, because all of the reactions and the subsequent washing steps in the tips are conducted by repeated aspiration and dispensing of reagents (repeated more than 100 times), the assay would be difficult to perform manually, and an automated liquid handling system would be necessary to avoid tedious manual pipetting and ensure the assay's reproducibility, which limits the application of this technology in most biological laboratories.

## 3. Conclusions

The assay performance of each technology achieved to date is summarized in Table 1. Most affinity-MS assays have achieved LODs of femtomole to attomole levels per analyzed sample such as plasma, corresponding to low ng/mL or high pg/mL concentrations. The iMALDI assay targeting EGFR detected 1 amol protein in buffer and a slightly higher amount in a few human breast cancer cells. The assay's LOD depends strongly on the quality of the antibodies and the sequence of the target peptides. In terms of reproducibility, most methods have achieved CVs of less than 15%, meeting the requirements in the FDA guideline for bioanalytical method validation [63]. There were relatively large quantitative variations for the mass-tag methods, but these could be improved by further optimizing tagging reactions and other assay conditions. Automation of liquid handling procedures generally improves assay reproducibility. Furthermore, it has been demonstrated that targeted immuno-MS assays can be highly multiplexed, as shown by a 172-plex tip-based immuno-MS approach. The cost and time required for anti-peptide antibody production, however, is still a major challenge in multiplexing these assays. The sample volumes required for each assay are comparable, and it has been observed that increasing the sample volume generally improves the LLOD. Lastly, high assay throughput is important for screening a large number of samples, which is a critical step for biomarker validation due to the large variations across populations [64, 65]. Assay automation has enabled the screening of more than 1000 samples per day, making it suitable for large-scale screening assays. These results demonstrate that affinity-MS could be a robust analytical tool for verification and validation studies of potential protein biomarkers of various diseases.

**Table 1.** Comparison of the assay performance using different immuno-enrichment techniques and ionization methods

| Immuno-enrichment techniques | LOD | CV | Multiplexing-capability achieved | Required sample volume | Assay throughput achieved |
| --- | --- | --- | --- | --- | --- |
| SELDI | < 10 fmol using 150 kD protein spots [37] | <11% [43] | 5-plex [38] | 50-150 μL [38] | 125 samples/assay [38] |
| SPR-immuno MALDI-TOF | Femtomole for LAG3 in human plasma [46] | Not available | 4-plex [45] | 0.5 μL [45] | Not available |
| Mass-tag MALDI | 200 pg/mL (60 amol/spot) for PSA [50] | Large, depending on efficiency of tagging reaction [50] | Single-plex | 10 μL [50] | Not available |
| iMALDI | 1 amol in buffer, EGFR from a few cells [10] | < 10% [15] | 2-plex [14] | 30 μL [15] | 188 samples/assay [15] |
| SISCAPA | Low pg/ml in plasma (amol range) [18] | 3.4% intra-run<br>4.3% inter-run [20] | 69-plex [25] | 30 μL [62] | 1500 samples in ~ 4 h [78] |
| In 96-well plate (ELIMSA) | 50 pg/mL PSA from 100 μL plasma (192 amol) [53] | <10% [79] | Single-plex | 50-100 μL [53] | 96-well plate based |
| In column | 0.5 fmol peptides in urine [54] | <15% [54] | Single-plex | 25-100 μL [54] | A hundred samples within 24 h [55] |
| In tips | 1.5 ng/mL for IGF 1 (7.8 fmol) [61] | < 10% intra- and inter-day [61] | 172-plex [62] | 40 μL plasma [61] | 1054 samples per day [61] |

## 4. Future perspectives

Antibody production still constitutes a major bottleneck in the development of affinity-MS assays. On one hand, affinity-MS only needs one antibody for each target, without the need to generate two antibodies against different epitopes. In addition, antibody selectivity is a less stringent requirement due to the second round of selection by mass spectrometry [4]. Also, for bottom-up assays the production of anti-peptide antibodies is more straightforward than those against intact proteins, because these peptides can be synthesized with high purity using an automated synthesizer [66]. On the other hand, affinity-enrichment of peptides is still restricted by the number of available antibodies, because most antibodies in the market target intact proteins and not peptides, thereby limiting the large-scale quantification of proteins based on the enrichment of their proteotypic peptides. To achieve proteome wide screening, more than 10,000 antibodies would have to be generated for affinity-MS [67], with a cost of approximately $500 for each antibody.

Several strategies have shown promise for tackling the antibody production challenge for bottom-up affinity-MS. If immuno-enrichment can be performed at the protein level, the antibodies targeting intact proteins can be applied, significantly expanding the availability of the usable antibodies. The captured intact proteins can be eluted for digestion [68], or can be digested directly on beads, on chips or in tips, followed by elution and ionization [46]. In addition, affinity binders capable of targeting multiple peptides per binder (named the Global Proteome Survey and Triple X Proteomics) have been developed as emerging approaches for producing antibodies at lower cost [66]. Alternatively, synthetic aptamers provide another option complementary to antibodies, and have shown lower background and easier production than antibodies [69].

While immuno-enrichment combined with MS-based assays could improve LODs by as much as 8,000fold [70], and have brought the assay LODs from the μg/mL range down to the low ng/mL range, most current affinity-MS assays are still 3-4 orders of magnitude less sensitive than the well-developed ELISA technology, which has achieved LODs in the low pg/mL range [71]. A low LOD is particularly important for early diagnosis of complex diseases such cancer. It has been estimated that with current protein quantification techniques in blood, a tumor might not be detected until it reaches the diameter of tens of millimeters, ten years after the initiation of the tumor [72]. To reach lower LODs, improved mass spectrometers might be necessary for better ionization and detection sensitivities.

A promising trend in the technology development would be to integrate affinity-MS with microfluidic chips for automated and simplified sample preparation and for reducing reagent costs [73, 74]. More importantly, through miniaturization the protein or peptide signals could be concentrated in smaller areas (in MALDI) or volumes (in ESI) therefore improving assay sensitivity [75, 76].

In terms of applications, affinity-MS is a powerful tool for the study of protein PTMs. Both modified and non-modified intact peptide isoforms can in theory be measured with a single antibody, as long as the target epitope falls within a region that is known to not be modified, and can be distinguished by a mass spectrometer. However, it is still challenging to measure a modification-containing peptide that is not proteotypic [25]. Another challenge is the analysis of low-abundance multiply-modified peptides, each of which generates a signal at a different m/z, thereby each variant could potentially fall below the LOD.

DBS appears the most favourable technique to date for personalized medicine and wellness. For regular health check-ups or for follow-up tests of chronic diseases, patients could easily collect DBS samples on a collection card by themselves, put the card in an envelope, and mail it to a centralized laboratory [65]. Personalized diagnosis can be performed by comparing new data with each individual's own baseline instead of with a cut-off based on population-wide averages. In DBS, however, not all proteins can be preserved well at the protein level, but most proteins are probably detectable at the peptide level after drying and long-term storage [77]. Affinity-MS has been recently used for quantifying peptides from DBS [27]. We can expect that more efforts will be made on time-course studies of peptides from DBS using affinity-MS, and these efforts will finally contribute to personalized management of diseases.

Finally, the gap between discovery and validation needs to be bridged by inexpensive high-throughput methods in order to bring more biomarker studies into routine clinical practice.

## Acknowledgements

HL thanks a postdoctoral fellowship from the National Science and Engineering Research Council of Canada (NSERC). RP thanks the University of Victoria for financial support during his graduate studies. We are grateful to the Genomic Innovations Network from Genome Canada and Genome British Columbia (project codes 204PRO for operations and 214PRO for technology development) for technology development funding. C.H.B. is also grateful for support from the Leading Edge Endowment Fund.

